# Rethinking asexuality: the enigmatic case of functional sexual genes in *Lepraria* (Stereocaulaceae)

**DOI:** 10.1101/2024.06.11.598483

**Authors:** Meredith M. Doellman, Yukun Sun, Alejandrina Barcenas-Peña, H. Thorsten Lumbsch, Felix Grewe

## Abstract

**Background:** The ubiquity of sex across eukaryotes, given its high costs, strongly suggests it is evolutionarily advantageous. Asexual lineages can avoid, for example, the risks and energetic costs of recombination, but suffer short-term reductions in adaptive potential and long-term damage to genome integrity. Despite these costs, lichenized fungi have frequently evolved asexual reproduction, likely because it allows the retention of symbiotic algae across generations. The relatively speciose lichenized fungal genus *Lepraria* is thought to be exclusively asexual, while its sister genus *Stereocaulon* completes a sexual reproductive cycle. A comparison of sister sexual and asexual clades should shed light on the evolution of asexuality in lichens in general, as well as the apparent long-term maintenance of asexuality in *Lepraria*, specifically.

**Results:** In this study, we assembled and annotated representative long-read genomes from the putatively asexual *Lepraria* genus and its sexual sister genus *Stereocaulon*, and added short-read assemblies from an additional 22 individuals across both genera. Comparative genomic analyses revealed that both genera were heterothallic, with intact mating-type loci of both idiomorphs present across each genus. Additionally, we identified and assessed 29 genes involved in meiosis and mitosis and 45 genes that contribute to formation of fungal sexual reproductive structures (ascomata). All genes were present and appeared functional in nearly all *Lepraria*, and we failed to identify a general pattern of relaxation of selection on these genes across the *Lepraria* lineage. Together, these results suggest that *Lepraria* may be capable of sexual reproduction, including mate recognition, meiosis, and production of ascomata.

**Conclusions:** Despite apparent maintenance of machinery essential for fungal sex, over 200 years of careful observations by lichenologists have produced no evidence of canonical sexual reproduction in *Lepraria*. We suggest that *Lepraria* may have instead evolved a form of parasexual reproduction, perhaps by repurposing *MAT* and meiosis-specific genes. This may, in turn, allow these lichenized fungi to avoid long-term consequences of asexuality, while maintaining the benefit of an unbroken bond with their algal symbionts.

## Background

Sexual reproduction is widespread among eukaryotes, suggesting its critical evolutionary advantage. It is widely recognized that sexual recombination promotes novel trait combinations and facilitates the purging of harmful alleles, thereby driving important adaptation [1–4]. However, the prevalence of obligate sexuality remains an intriguing conundrum due to its numerous costs, including the breakup of favorable gene combinations, the risk and energy requirements associated with recombination, and the costs of finding a mate (reviewed in [5]). Conversely, an asexual individual avoids these costs and can maintain beneficial adaptations in stable environments. However, asexuality also has long-term costs, primarily the decreased effectiveness of selection due to the complete linkage of genetic variants, which reduces adaptive potential and the ability to maintain variation in the presence of strong fluctuating selection [6]. Over time, asexual lineages may also suffer declines in the integrity and functionality of the nuclear genome, as evidenced by accumulating genome damage, such as loss of telomeres [7], indirect effects of mitochondrial metabolism [8, 9], and proliferation of transposable elements [10]. Thus, while asexuality may circumvent the immediate costs of sexual reproduction, it introduces significant risks to genomic health over time and potentially compromises an organism’s ability to adapt and survive in changing environments.

Despite the long-held belief in exclusively asexual reproduction among certain eukaryotic lineages, recent studies increasingly reveal evidence of cryptic sex, or at least recombination, challenging these assumptions [11–14]. The genomic revolution has led the way in changing this paradigm; the remarkable conservation of basic cellular processes and genes controlling meiosis across eukaryotes [15] has been broadly leveraged for the bioinformatic identification of sex and meiosis machinery across many putatively asexual lineages [13]. For instance, numerous fungi once labeled as “*fungi imperfecti*” or “Deuteromycota” have been shown to harbor functional mating-type and meiosis genes, demonstrating their capacity for sexual reproduction (reviewed in [5]). In a recent case, the genus *Pseudozyma* was studied, and it was found that species previously thought to be asexual have functional mating and meiosis genes in their genomes [16]. Similarly, in *Verticillium dahliae*, a species traditionally considered to reproduce only asexually, the presence of mating and meiosis genes, as well as recombination, was confirmed through genotyping-by-sequencing [17]. These findings reveal a previously unrecognized complexity in fungal reproductive strategies, potentially impacting our understanding of their ecological adaptability and evolutionary dynamics.

Lichenized fungi, growing in symbiosis with an algal and/or cyanobacterial partner, commonly exhibit sexual and asexual reproduction (reviewed in [18]). Sexual lichens undergo mating and meiosis, form reproductive structures primarily called ascomata, and release fungal spores. In addition to conventional costs of sexuality, these fungal spores must regain their algal symbiont to form the lichen thallus. Sexual reproduction in lichenized fungi is regulated by genes located at the mating type locus (*MAT*), which includes two distinct allelic variants: the *MAT1-1* and *MAT1-2* idiomorphs. Lichenized fungi can be either homothallic with both idiomorphs encoded in the same genome, where a single thallus can self-fertilize, or heterothallic, requiring the interaction between genetically distinct thalli, each carrying a different idiomorph [19–23]. On the other hand, asexual lichens maintain an association between the fungus and the algal symbiont throughout reproduction, forming asexual propagules (soredia, isidia, etc.), small fragments of thallus incorporating the full lichen community (fungus, algae, bacteria, etc.) [24]. The lichenized fungal genus *Lepraria* is thought to be exclusively asexual [25]. Lichenized fungi form stable and usually long-lived vegetative structures and ascomata in these fungi usually remain for long periods of time, mostly several years [26]. The formation of meiospores, which are formed within ascomata in filamentous ascomycetes, has never been observed among members of *Lepraria*, and its morphology is composed primarily of soredia [27]. However, despite this apparent asexuality, *Lepraria* is a relatively speciose lineage [25, 27–29]. Conversely, its sister genus *Stereocaulon* reproduces sexually [25, 30]; therefore, comparative genomic analysis has the potential to highlight the long-term genomic effects of asexuality. For example, under a “use it or lose it” model, a truly asexual lineage should experience relaxed selection on unused reproductive genes, leading to an accumulation of deleterious mutations, pseudogenization, and gene loss within a few million years [31]. Thus, we might expect to see a loss or degradation of genes involved in mate recognition, meiosis, and/or the development of reproductive structures across *Lepraria* [32].

We address the apparent asexuality of *Lepraria* using a three-pronged comparative genomic approach. First, we ask if *Lepraria* have functional mating systems by identifying *MAT* loci and discerning idiomorphs for *Lepraria* and *Stereocaulon* samples. Second, we ask if *Lepraria* is capable of performing meiosis and/or developing ascomata by identifying and assessing the functionality of core meiotic and fruiting body developmental genes. Third, we ask if these core genes are likely experiencing ongoing selection by testing for evidence of relaxation of selection in the putatively asexual *Lepraria* lineage. By comparing sister sexual and asexual clades, we hope to gain insights into the evolution of asexuality in lichens and probe whether *Lepraria* is a truly asexual lineage.

## Methods

### Sample collection, DNA extraction, and sequencing for reference genomes

*Lepraria finkii* was collected from Chicago, IL, in June 2017, and *Stereocaulon virgatum* was collected from Antarctica in November 2017 (Table 1). Samples were identified using morphological characteristics and resolution of secondary metabolites with high-performance thin-layer chromatography with solvent C, following established methods [33–36]. For the sexual *S. virgatum*, axenic cultures were produced from ascospores and grown on malt-yeast extract agar until the individual culture reached sufficient size for DNA extraction. As *L. finkii* does not produce ascospores, a fungal culture could not be isolated, and the entire symbiotic organism was processed.

**Table 1:** Sample collection information and genome assembly statistics.

| Species | Size (Mb) | Scaffolds | N50 | BUSCO | Location | Date | Lat. | Long. | Source |
| --- | --- | --- | --- | --- | --- | --- | --- | --- | --- |
| *Lepraria caerulescens* | 37.7 | 8179 | 15568 | 96.9% | Antarctica | 05-Mar-18 | -62.66 | -60.39 | This study |
| *Lepraria chileana* | 37.6 | 9474 | 21804 | 95.7% | Chile | 19-Dec-17 | -42.62 | -74.11 | This study |
| *Lepraria eburnea* | 34.6 | 8226 | 16078 | 97.3% | New Zealand | 03-Dec-17 | -45.91 | 169.49 | This study |
| *Lepraria finkii* (1) | 41.4 | 304 | 402087 | 96.5% | United States | 21-Jun-17 | 42.15 | -87.79 | This study |
| *Lepraria finkii* (2) | 46 | 9450 | 37458 | 97.7% | Chile | 17-Dec-17 | -41.59 | -72.59 | This study |
| *Lepraria lobificans* | 43.3 | 4750 | 49780 | 97.1% | New Zealand | 18-Nov-17 | -45.90 | 170.48 | This study |
| *Lepraria neglecta* | 42 | 11 | 5201525 | 97.4% | United States | 25-Sep-19 | 47.48 | -117.56 | GCA_033220425.1 |
| *Lepraria neozelandica* | 36.6 | 8490 | 13080 | 97.1% | New Zealand | 06-Nov-17 | -36.80 | 174.70 | This study |
| *Lepraria nothofagi* | 32.5 | 5143 | 25114 | 97.6% | Antarctica | 24-Feb-28 | -68.66 | -60.40 | This study |
| *Lepraria sp. 1* | 27.3 | 11508 | 3875 | 84.1% | New Zealand | 26-Aug-17 | -45.85 | 170.67 | This study |
| *Lepraria sp. 2* | 36.3 | 6137 | 27309 | 97.4% | Taiwan | 07-Nov-19 | 24.22 | 121.32 | This study |
| *Lepraria straminea* | 29.2 | 3121 | 24536 | 97.0% | Antarctica | 25-Feb-18 | -62.66 | -60.39 | This study |
| *Lepraria toilenae* | 28.9 | 5542 | 12595 | 96.1% | Australia | 25-Jan-16 | -38.42 | 146.72 | This study |
| *Stereocaulon alpinum* (1) | 38.3 | 546 | 289189 | 97.6% | Antarctica | 04-Feb-06 | -62.13 | -58.46 | GCA_018257865.1 |
| *Stereocaulon alpinum* (2) | 26.2 | 4475 | 13299 | 96.5% | Antarctica | 24-Feb-18 | -68.66 | -60.40 | This study |
| *Stereocaulon antarcticum* | 23.9 | 6706 | 7311 | 95.8% | Antarctica | 22-Feb-18 | -68.66 | -60.40 | This study |
| *Stereocaulon caespitosum* | 22.2 | 6146 | 6497 | 93.8% | New Zealand | 01-Dec-17 | -45.83 | 170.46 | This study |
| *Stereocaulon cf. fronduliferum* | 17.4 | 9503 | 2888 | 80.2% | New Zealand | 01-Dec-17 | -45.83 | 170.46 | This study |
| *Stereocaulon cornutum* | 26.8 | 5699 | 13056 | 97.0% | Cuba | 09-Jul-16 | 20.01 | -76.84 | This study |
| *Stereocaulon glabrum* (1) | 27 | 5277 | 13986 | 97.0% | Chile | 13-Dec-17 | -55.00 | -67.68 | This study |
| *Stereocaulon glabrum* (2) | 26.1 | 5103 | 12704 | 96.8% | Antarctica | 06-Mar-18 | -62.64 | -60.37 | This study |
| *Stereocaulon gregarium* | 16.3 | 6687 | 3451 | 83.6% | New Zealand | 16-Nov-17 | -45.81 | 170.55 | This study |
| *Stereocaulon vesuvianum* * | 27 | 3401 | 20864 | 97.3% | Denmark | 02-Jul-19 | 55.87 | 9.83 | This study |
| *Stereocaulon virgatum* | 34.7 | 264 | 721891 | 97.3% | Dominica | 13-Aug-17 | 15.34 | 61.31 | This study |
| *Cladonia borealis* | 36 | 48 | 1657173 | 96.5% | Antarctica | 28-Jan-06 | -62.13 | -58.47 | GCA_018257855.2 |
| *Cladonia squamosa* | 38.1 | 121 | 1635907 | 97.3% | United Kingdom | 02-Apr-21 | 56.02 | 56.02 | GCA_947623385.2 |

\* *Stereocaulon vesuvianum* var. *symphycheiloides*

High-molecular-weight DNA extractions of the *S. virgatum* fungal culture and *L. finkii* sample were based on an existing protocol, with some modifications [37]. About 0.6 g of dried material was flash-frozen with liquid nitrogen, ground with a ceramic mortar and pestle, and then allowed to reach room temperature. The ground material was incubated with 500 μL lysis buffer and 20 μL proteinase K at 64 °C up to 4 h, then cooled on ice for 5 min. To the cool mixture, 100 μL of 5 M KAc was added and incubated for 5 min on ice, then centrifuged at max speed at 4 °C for 10 min. The supernatant was added to 500 μL phenol:chloroform:isoamyl alcohol and centrifuged at max speed at 4 °C for 10 min. The supernatant was then added to 500 μL isopropanol and cooled at −80 °C for 1 h. The isolated DNA was pelleted at maximum speed at 4 °C for 30 min, washed twice with 1 mL 70% ethanol, and eluted in 50 μL TE buffer.

Extracted DNA samples were submitted to the DNA services facility of the University of Illinois at Urbana-Champaign for library construction and sequencing. For *L. finkii,* genomic DNA was sheared in a gTube (Covaris, Woburn, MA) for 1 min at 6,000 rpm in an Eppendorf MiniSpin plus microcentrifuge (Eppendorf, Hauppauge, NY). The sheared DNA from each sample was converted into an Oxford Nanopore (ON) library with the 1D library kit SQK-LSK108 and sequenced on a SpotON R9.4.1 FLO-MIN107 flowcell for 48hs, using a GridION X5 sequencer (Oxford Nanopore Technologies, Oxford, UK). Basecalling was performed with Guppy 1.8.5-1, resulting in approximately 3.7 Gb of data across 942,752 reads, averaging 3.9 kb per read. The *S. virgatum* library was prepared according to low input protocol modifications. After shearing for 1 min at 5,000 rpm, the sheared DNA was size selected on a 1% agarose gel. Whole genome PCR amplification was performed with 200ng of the excised DNA. Then 925ng of PCR-amplified and 590ng of native DNA were mixed, converted into an ON library, and sequenced as above. Basecalling was performed with MinKNOW 2.0, resulting in approximately 1.3 million reads, averaging 7.5 kb in length and totaling ∼9.6 Gb [38]. Both DNA samples were also converted into Illumina sequencing libraries with the KAPA HyperPrep library construction kit (Roche Sequencing Solutions, Indianapolis, IN) and paired-end sequenced for 150 cycles on a NovaSeq6000 (Illumina, San Diego, CA) [38].

### Reference genome assembly

Raw ON reads for both *L. finkii* and *S. virgatum* were trimmed with Porechop v0.2.4 to remove adaptors and filtered with Filtlong v0.2.1 to maintain the best 90% of bases, with a minimum read length of 1000 bp [39]. Illumina reads were quality checked with FastQC v0.12.1 [40] and processed with Trimmomatic v0.39-2 [41], using the following parameters: *ILLUMINACLIP:TruSeq3-PE.fa:2:30:10 LEADING:10 TRAILING:10 SLIDINGWINDOW:4:15 MINLEN:25*. Filtered ON reads were assembled with Flye v2.8, with the *--nano-raw* flag and an estimated genome size of 45 Mb for *S. virgatum* and 150 Mb for *L. finkii*, to account for the photobiont and potential bacterial symbiont genomes [42]. This assembly was then polished with two rounds of Racon v1.4.3 [43] and one round of Medaka v1.6.0 [44], followed by four rounds of polishing with the filtered Illumina reads using NextPolish v1.4.0 [45], all with default parameter settings. Fungal contigs were identified from the *L. finkii* metagenome assembly using Kraken2 v2.1.2 and a custom database constructed with the pre-built *fungi* library and our *S. virgatum* reference genome [46]. Genome contiguity was assessed with QUAST v5.2.0 [47] and completeness was evaluated with BUSCO v5.4.7 [48], using the dataset for Ascomycota (ascomycota_odb10).

### RNA extraction, sequencing, and reference genome annotation

*Lepraria finkii* RNA was extracted from a thallus fragment with the Norgen Plant/Fungi Total RNA purification kit (Norgen Biotek, Thorold, Ontario, Canada). The tissue was homogenized for 1.5 minutes in 600 uL of Lysis Buffer C, using ClaremontBio microHomogenizer 2.0 mL tubes (Claremont BioSolutions LLC, Upland, CA). Extraction then proceeded per the Norgen kit protocol. Stranded poly(a) selected libraries were constructed with the Tecan mRNA-Seq for MagicPrep NGS kit (Tecan Life Sciences, Männedorf, Switzerland); 50 ng of RNA were loaded into the Tecan MagicPrep™ NGS automated library preparation system and 18 PCR cycles were selected, according to protocol recommendations.

*Stereocaulon virgatum* RNA was extracted from the fungal culture using the Maxwell® RSC Plant RNA Kit (Promega, Madison, WI). Tissue was homogenized in 30 second intervals for 1.5 minutes at 30hz, using 2mm ZR BashingBead Lysis Tubes (Zymo Research, Irvine, CA) in the Qiagen TissueLyser II (Qiagen, Hilden, Germany). The extraction was completed according to the Maxwell® RSC 48 Instrument (Promega, Madison, WI) protocol. Stranded poly(a) selected libraries were then prepared with the NEXTFLEX® Rapid Directional RNA-Seq Kit 2.0 (PerkinElmer Inc., Waltham, MA). For both species, 150 bp paired-end sequences were generated on the NovaSeq6000 (Illumina, San Diego, CA) platform [38].

For each genome, we used RepeatModeler v2.0.4 to generate a species-specific repeat library, which we combined with the last available version of the RM RepBase library (RepBaseRepeatMaskerEdition-20181026) for masking [49]. With RepeatMasker v4.1.5 [50], we sequentially masked the simple repeats, fungal elements sourced from RepBase, followed by the known and then unknown repeats, following the suggestion of Koochekian et al. [51]. Each combined soft masked genome was then annotated with *funannotate* v1.8.15 [52]. Briefly, *funannotate* was trained with the respective RNA-seq data set, predictions were made with default settings, and UTRs were updated with the RNA-seq data set. The evidence modeler settings were left as defaults, resulting in 11,218 gene models supported by 12,128 total transcripts for *S. virgatum* and 14,168 gene models supported by 15,107 transcripts for *L. finkii*.

### Sample collection, DNA extraction, and sequencing for metagenomes

Collection details for an additional 11 *Lepraria* and 9 *Stereocaulon* samples are included in Table 1. Samples were identified to species level where possible, as above, using morphology and thin-layer chromatography [33–36]. Total genomic DNA was extracted from thallus fragments using the ZR Fungal/Bacterial DNA Miniprep Kit (Zymo Research Corp., Irvine, CA, USA), following the manufacturer’s instructions. DNA concentration was verified using the Qubit® dsDNA HS Assay Kit (Life Technologies, Grand Island, NY), and DNA was submitted to the University of Wisconsin-Madison Biotechnology Center. Samples were prepared according to the Celero PCR Workflow with Enzymatic Fragmentation (Tecan Genomics, Redwood City, CA). Quality and quantity of the finished libraries were assessed using an Agilent Tapestation (Agilent, Santa Clara, CA) and Qubit® dsDNA HS Assay Kit, respectively. Paired-end, 150 bp sequencing was performed using the Illumina NovaSeq6000 (Illumina, San Diego, CA) [38].

### Metagenome assembly and annotation

Quality of the Illumina sequences was assessed with FastQC, and they were trimmed to remove adapter contamination and low-quality bases with Trimmomatic, as above [40, 41]. Kraken2 was used to filter trimmed reads, based on a database constructed from the representative *L. finkii* and *S. virgatum* genomes [46]. We assembled filtered reads with the *--careful* option in Spades v3.15.5 [53], and contigs less than 500bp in length were removed with seqkit v2.5.1 [54]. Assemblies were then annotated with *funannotate*, using *Aspergillus nidulans* as the BUSCO and Augustus seed species, with remaining default settings [52]. Finally, we assessed genome contiguity and completeness with QUAST and BUSCO, as above [47, 48].

### Phylogeny

Genome assemblies for *Cladonia borealis* and *C. squamosa* were downloaded from NCBI to use as outgroups (GCA_018257855.2 and GCA_947623385.2, respectively), and *L. neglecta* and *S. alpinum* were added to the main data set (GCA_033220425.1 and GCA_018257865.1, respectively). Associated annotations were downloaded for *C. borealis* and *L. neglecta*, while *funannotate* was used for *C. squamosa* and *S. alpinum* gene prediction, following the methods detailed for the metagenomes above [52]. Based on predicted proteins from annotations of all included *Lepraria*, *Stereocaulon*, and *Cladonia* (Table 1), OrthoFinder v2.5.5 identified single-copy orthologs present in all 26 taxa [55]. A concatenated multiple sequence alignment of these proteins was generated with the MAFFT option in OrthoFinder. RAxML v8.2.12 was run with the PROTGAMMAAUTO option, which chose the JTT likelihood model, and 500 rapid bootstraps [56]. The final tree was edited in FigTree v1.4.4 [57].

### MAT locus assembly and annotation

The *MAT* loci in the reference *L. finkii* and *S. virgatum* were identified via a *diamond blastx* v2.1.6 search for the conserved flanking genes *apn2* and *sla2*, using *Saccharomyces cerevisiae* proteins [58]. The idiomorphs were identified by *blastp* search of the predicted annotations between these two genes, as well as by comparison of the gene orientation and structure with published Lecanoromycetes *MAT* loci [22, 59]. We identified the *MAT* locus from the remaining *Lepraria* and *Stereocaulon* genomes by mapping the *apn2* and *sla2* sequences extracted from their respective reference genomes to each additional genome in Geneious Prime v2023.1.2. Unique localization of the *MAT* locus in each genome was confirmed by *blastx* search. As above, we assigned an idiomorph based on the structure and orientation of the predicted genes within the *MAT* region and confirmed sequence similarity via alignment with the Clustal Omega v1.2.3 progressive alignment algorithm in Geneious.

### Meiosis and ascomata developmental gene functionality

Meiosis toolkit genes were collated from bioinformatic analyses of putatively asexual genomes [13, 15, 16, 60, 61]. We downloaded a representative fungal protein sequence for each gene from Uniprot or NCBI, created a diamond database, and identified the best match in the *L. finkii* and *S. virgatum* reference genomes using *diamond blastx*. The regions containing the matching annotations and flanking genes were extracted from each genome and aligned to examine synteny and compare gene models. In addition, we examined the mapping of each assembled transcriptome to its respective genome, generated by *funannotate*. For each focal gene and genome, we then compared the predicted gene model to the mapped transcripts, confirming intron boundaries and start and stop codons. Finally, we compared predicted models and transcripts across *L. finkii* and *S. virgatum*, and manually curated a consensus gene model, if necessary. Many meiosis and mitosis genes are members of gene families that evolved from extensive duplications of DNA repair genes (e.g., *dmc1* and *rad51*; *rec8* and *rad21*; *pms1* and *mlh1/2/3*; *msh2/4/5/6*; *mnd1* and *hop1/2*) [13]. To confirm the analysis of the correct paralog, we extracted the translated protein sequence of each gene in the family from the *L. finkii* and *S. virgatum* annotations, aligned them with the representative Uniprot/NCBI fungal proteins with MUSCLE v5.1, and generated neighbor-joining trees with Geneious Tree Builder. Ascomata development genes were identified from two reviews on the formation of sexual structures in model fungi [62, 63]. Orthologs were identified and extracted from the *L. finkii* and *S. virgatum* reference genomes following the methods detailed above for the meiosis toolkit genes.

### RELAX analysis

Target genes identified as part of the meiosis toolkit and putatively involved in the development of ascomata were identified in all additional assemblies by mapping each gene extracted from the *L. finkii* and *S. virgatum* reference genomes to each remaining assembly in Geneious Prime. They were then extracted with up and downstream flanking genes, if assembly contiguity allowed, and these regions were aligned with the Clustal Omega algorithm. Annotated coding sequence predictions were manually adjusted in Geneious, based on the *L. finkii* and *S. virgatum* gene models, and discarded if the assembly of the target gene was incomplete. The coding sequences were exported in fasta format, and a codon-aware alignment was performed with MACSE v2.07 [64]. To match each fasta, the OrthoFinder species tree was pruned using the keep.tip function in ape v5.7.1 (R version 4.3.2) to remove taxa with fragmented genes [65]. *Lepraria* nodes were labeled as test (T), *Stereocaulon* as reference (R), and outgroup as nuisance with the online tool phylotree.js. RELAX was run with the codon-aware alignment, labeled tree, and default settings using the command line implementation of HYPHY v2.5.48 [66, 67].

## Results

### Genome sequencing of Lepraria and Stereocaulon

In this study, we assembled one representative high-quality ON genome for each of the sister genera *Lepraria* and *Stereocaulon* (Table 1). The *L. finkii* assembly consisted of 41.2 Mb, across 304 scaffolds, with an N50 of approximately 402 kb and 96.5% BUSCO completeness.

*Stereocaulon virgatum* was 34.7 Mb, on 264 scaffolds, with an N50 of 722 kb and was 97.3% complete. An additional 11 *Lepraria* and 9 *Stereocaulon* genomes were assembled from metagenomic Illumina data filtered for fungal reads. For these fungal genomes, genome size ranged from 16.3 Mb for *S. gregarium* to 46 Mb for *L. finkii*, with a median of 28.1 Mb. The median scaffold number was 6141.5, ranging from 3121 in *L. straminea* to 11,508 in *Lepraria sp. 1*. Scaffold N50s were between 2.9 (*S. cf. fronduliferum*) and 49.8 kb (*L. lobificans*), with a median of 13.6 kb, while BUSCO completeness had a median of 96.95% and spanned from 80.2% in *S. cf. fronduliferum* to 97.7% in *L. finkii* (Table 1, Fig. S1).

Analysis of predicted genes with Orthofinder resulted in 1262 single-copy orthologs present in all 26 taxa. Rooting with an outgroup consisting of *Cladonia borealis* and *C. squamosa*, the maximum likelihood phylogenetic tree confirmed that *Lepraria* and *Stereocaulon* are reciprocally monophyletic clades (Fig. 1B). The tree was well-resolved, and all nodes had high bootstrap support (>90%).

**Figure 1:**
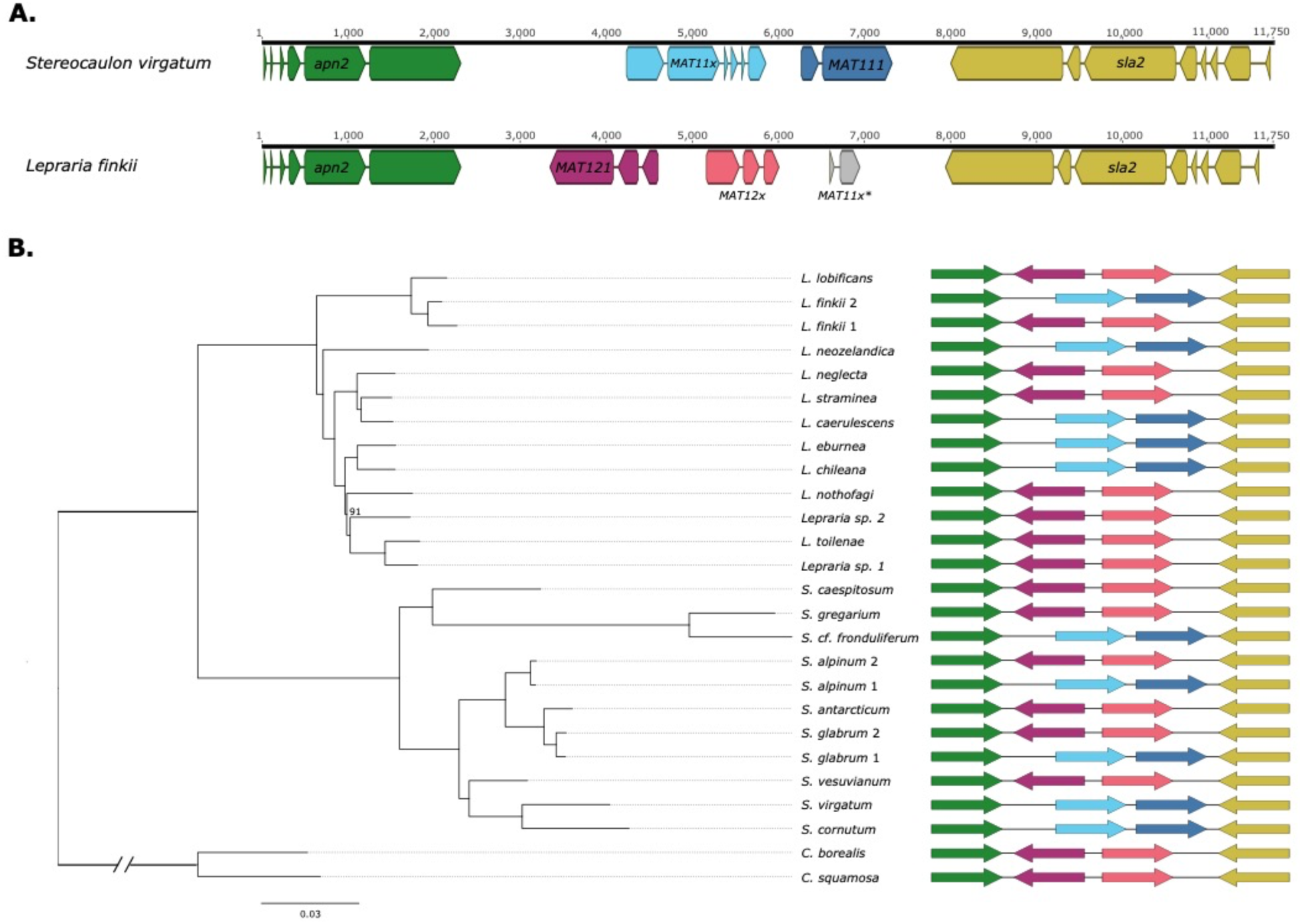
Mating-type locus identities and phylogenomic tree for *Stereocaulon* and *Lepraria* samples. A) Schematics representing the *MAT* locus gene annotations for the *Stereocaulon virgatum* (*MAT1-1* idiomorph) and *Lepraria finkii* (*MAT1-2* idiomorph) genomes. Cartoon arrows represent exons, with introns indicated by intervening lines. Using RNAseq data from each species, gene models were predicted with *funannotate* and curated by manual inspection of the mapped transcriptome. Gene identities were confirmed with *blastp* searches with predicted protein queries. Gene models were modified from figures produced by Geneious version 2023.1 created by Biomatters. B) Phylogenomic tree of *Stereocaulon* and *Lepraria* samples, indicating the *MAT* idiomorph identified in each genome. The maximum likelihood tree was constructed with RAxML, based on 1262 single copy orthologs identified by OrthoFinder; all bootstraps not indicated are 100%. Color and orientation of the gene models (arrows) in the *MAT1-1* and *MAT1-2* idiomorphs match the schematics in Fig. 1A.

### The mating-type locus appears functional in Lepraria

In both *Stereocaulon virgatum* and *Lepraria finkii* ON assemblies, we identified complete mating-type loci with apparently functional *MAT* genes (Fig. 1A). In both genomes, the loci were assembled as contiguous, flanked by the upstream *apn2* and downstream *sla2* genes. The reference *S. virgatum* genome contained a *MAT1-1-1* idiomorph, with the *MAT1-1-1* gene downstream of the *MAT1-1-x* accessory gene, transcribed in the same direction. We did not name the accessory *MAT* gene because it is unnecessary within the scope of our study and the nature of the nomenclature is controversial [68]. An NCBI *blastp* conserved domain search confirmed the presence of the expected alpha domain in *MAT1-1-1* (Fig. S2). The *L. finkii* reference harbored a *MAT1-2-1* idiomorph; this gene and the accessory gene *MAT1-2-x* were divergently transcribed, with *MAT1-2-1* closer to *apn2*. In between the accessory gene and *apn2*, there was also a small predicted coding sequence with some homology to the 3’ end of the *MAT1-1-1* gene. An HMG (high mobility group) domain within *MAT1-2-1* was confirmed with an NCBI conserved domain search (Fig. S2B). The remaining *Lepraria* and *Stereocaulon* genomes showed a mix of *MAT1-1-1* (n=9) and *MAT1-2-1* (n=13) idiomorphs across our samples (Fig. 1B). In both *Stereocaulon* and, more importantly, *Lepraria*, where we had two individuals of the same species (i.e., *L. finkii, S. alpinum, S. glabrum*), the *MAT* loci were of opposite idiomorphs, suggesting the possibility of sexual reproduction. Alignment of all assembled *MAT* loci suggests that the *MAT1-1-1-like* small open reading frame in the *MAT1-2-1* idiomorph may have originated from a crossover event between the two idiomorphs (Fig. S3A); although *Stereocaulon* annotations do not predict a gene in that region, *blastx* searches with these sequences supported similarity with *MAT1-1-1*.

### Meiosis-specific genes appear functional across Lepraria

The survey of meiosis toolkit genes [15, 16, 60, 61] revealed no evidence that any of these genes had lost function in *Lepraria*. All nine meiosis-specific genes (*dmc1*, *hop1*, *hop2*, *mer3*, *mnd1*, *msh4*, *msh5*, *rec8*, and *spo11*) were identified in our *L. finkii* reference genome (Fig. 2A). All genes maintained the start and stop codons present in *S. virgatum* and could be confirmed in our transcriptome data. The NCBI *blastp* functional domain search results showed that all but one gene maintained the same functional domains in *Lepraria* and the known sexual *Stereocaulon* (Fig. S4). Unexpectedly, the core meiosis gene *hop2* appears to be pseudogenized in *Stereocaulon*, with an internal stop codon at amino acid position 74 of 234. This stop falls within the TBPIP/Hop2 winged helix domain identified by *blastp* (residues 50-94) in *Lepraria* (Fig. S4), suggesting that *Stereocaulon* HOP2 proteins are not functional.

**Figure 2:**
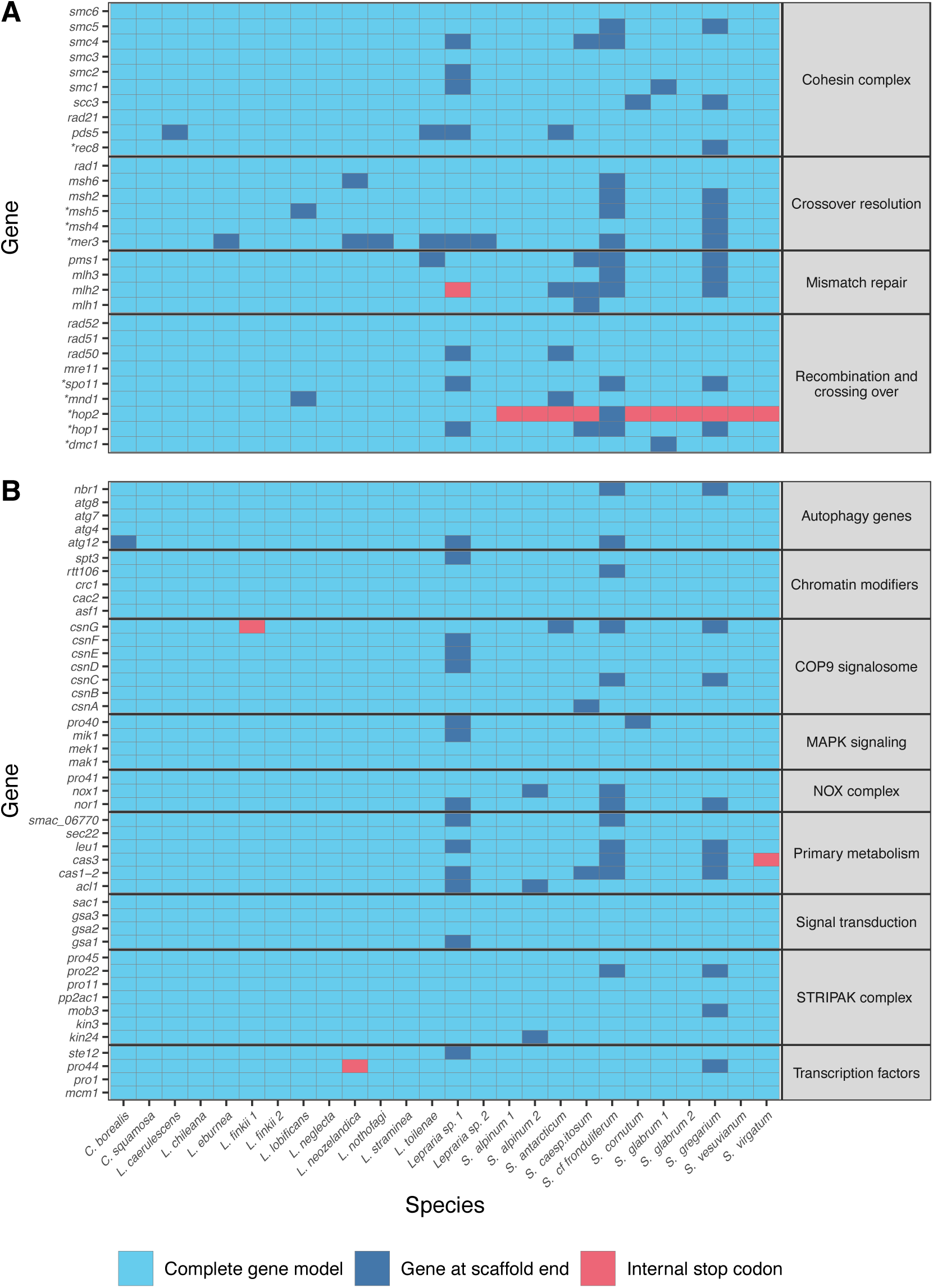
Presence and function of meiosis, mitosis, and sexual development genes across *Stereocaulon* and *Lepraria* samples. A) A heatmap displaying the presence of putatively functional meiosis and mitosis genes across *Stereocaulon* and *Lepraria*. The asterisks (*) indicate core meiosis-specific genes. With one major exception, all genes were present and complete, or partially present at the end of a scaffold. Interestingly, meiosis-specific *hop2* appears to have a premature internal stop codon in the sexual *Stereocaulon* lineage. B) A similar heatmap for ascomata development genes. No gene was missing completely, and all genes appeared generally functional across both *Stereocaulon* and *Lepraria* lineages.

These genes could also be extracted from the majority of the metagenome assemblies (Fig. 2A). For most genes, we were able to confirm conserved gene models across all *Lepraria* and *Stereocaulon* genomes. Across all genes, *S. cf. fronduliferum*, *S. gregarium*, and *Lepraria sp. 1* had the most fragmented genes at 12, 12, and 8, respectively. These were also the only three genome assemblies, with less than 90% BUSCO completeness (Table 1). Aside from *hop2* across all *Stereocaulon* samples, only *mlh2* in *Lepraria sp. 1*, a relatively low-quality assembly, appeared truncated. We identified no other premature stop codons in any species in any meiosis or mitosis gene.

Tests for relaxation of selection in *Lepraria* in the meiosis-specific genes compared to genes with mitotic functions also showed no specific pattern (Table 2). We found no evidence that relaxed selection was more common in any specific pathway, or among meiosis-specific genes compared to mitosis genes, as you would expect if these genes were truly free from selection in *Lepraria*.

**Table 2:**
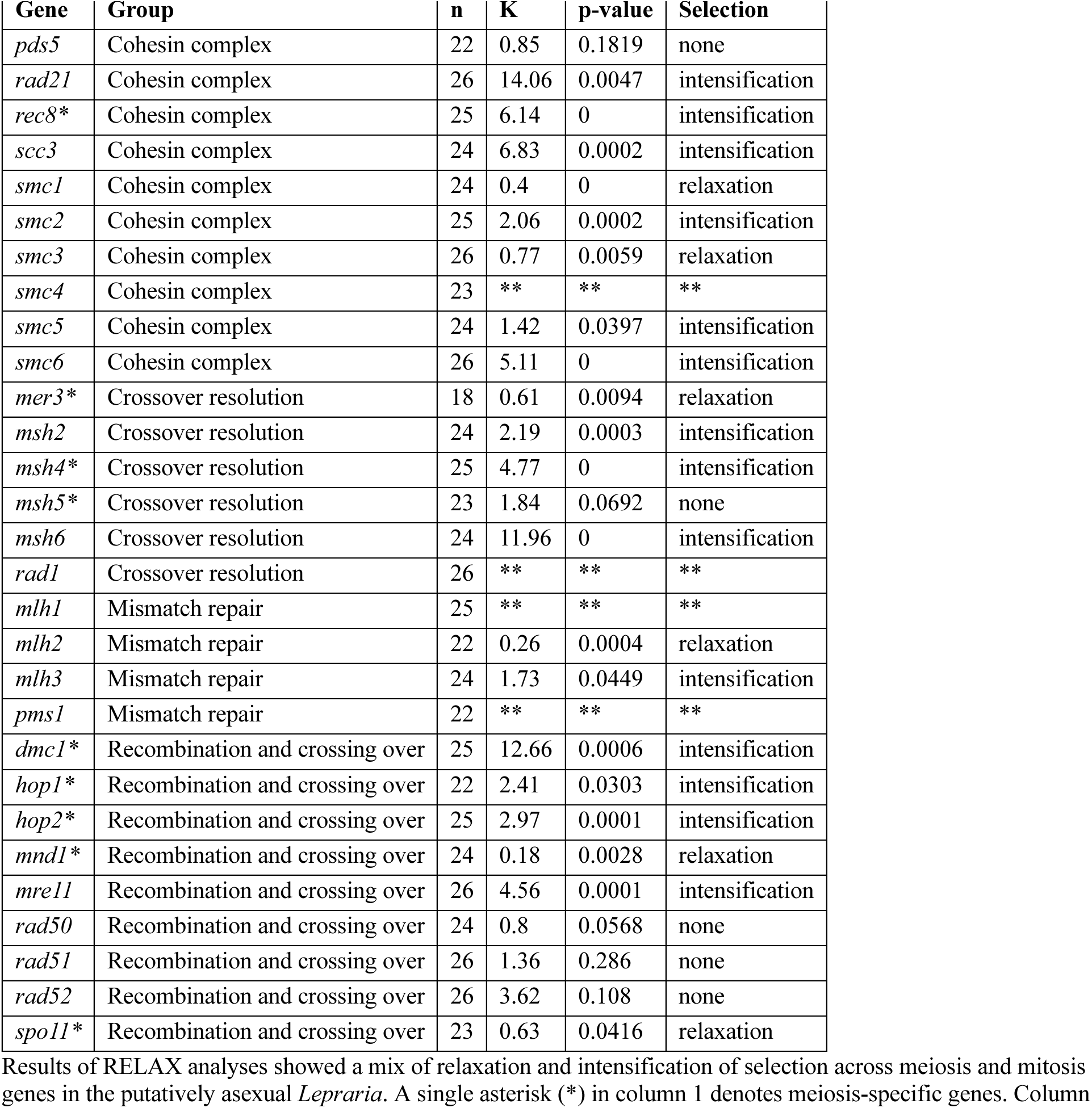
Selection on meiosis and mitosis genes in asexual *Lepraria* compared to sexual *Stereocaulon*.

| Gene | Group | n | K | p-value | Selection |
| --- | --- | --- | --- | --- | --- |
| *pds5* | Cohesin complex | 22 | 0.85 | 0.1819 | none |
| *rad21* | Cohesin complex | 26 | 14.06 | 0.0047 | intensification |
| *rec8\** | Cohesin complex | 25 | 6.14 | 0 | intensification |
| *scc3* | Cohesin complex | 24 | 6.83 | 0.0002 | intensification |
| *smc1* | Cohesin complex | 24 | 0.4 | 0 | relaxation |
| *smc2* | Cohesin complex | 25 | 2.06 | 0.0002 | intensification |
| *smc3* | Cohesin complex | 26 | 0.77 | 0.0059 | relaxation |
| *smc4* | Cohesin complex | 23 | ** | ** | ** |
| *smc5* | Cohesin complex | 24 | 1.42 | 0.0397 | intensification |
| *smc6* | Cohesin complex | 26 | 5.11 | 0 | intensification |
| *mer3\** | Crossover resolution | 18 | 0.61 | 0.0094 | relaxation |
| *msh2* | Crossover resolution | 24 | 2.19 | 0.0003 | intensification |
| *msh4\** | Crossover resolution | 25 | 4.77 | 0 | intensification |
| *msh5\** | Crossover resolution | 23 | 1.84 | 0.0692 | none |
| *msh6* | Crossover resolution | 24 | 11.96 | 0 | intensification |
| *rad1* | Crossover resolution | 26 | ** | ** | ** |
| *mlh1* | Mismatch repair | 25 | ** | ** | ** |
| *mlh2* | Mismatch repair | 22 | 0.26 | 0.0004 | relaxation |
| *mlh3* | Mismatch repair | 24 | 1.73 | 0.0449 | intensification |
| *pms1* | Mismatch repair | 22 | ** | ** | ** |
| *dmc1\** | Recombination and crossing over | 25 | 12.66 | 0.0006 | intensification |
| *hop1\** | Recombination and crossing over | 22 | 2.41 | 0.0303 | intensification |
| *hop2\** | Recombination and crossing over | 25 | 2.97 | 0.0001 | intensification |
| *mnd1\** | Recombination and crossing over | 24 | 0.18 | 0.0028 | relaxation |
| *mre11* | Recombination and crossing over | 26 | 4.56 | 0.0001 | intensification |
| *rad50* | Recombination and crossing over | 24 | 0.8 | 0.0568 | none |
| *rad51* | Recombination and crossing over | 26 | 1.36 | 0.286 | none |
| *rad52* | Recombination and crossing over | 26 | 3.62 | 0.108 | none |
| *spo11\** | Recombination and crossing over | 23 | 0.63 | 0.0416 | relaxation |

Results of RELAX analyses showed a mix of relaxation and intensification of selection across meiosis and mitosis genes in the putatively asexual *Lepraria*. A single asterisk (\*) in column 1 denotes meiosis-specific genes. Column

### Lepraria harbors genes for the development of ascomata

A survey of 45 genes involved in fruiting body formation in model fungi [62, 63] revealed that *Lepraria* likely has the ability to produce reproductive structures. Across the broad categories of autophagy genes, chromatin modifiers, MAPK signaling, primary metabolism, signal transduction, and transcription factors, as well as components of the COP9 signalsome and NOX and STRIPAK complexes, all genes were identified in the *L. finkii* reference genome. Aside from *csnG*, which appears to have an internal stop codon in the *L. finkii* reference, but not in any other *Lepraria* genome, all genes are present, with the same start and stop codons as in the *S. virgatum* reference genome. Across most of the 22 *Lepraria* and *Stereocaulon* metagenome assemblies, these genes were present and appeared functional (Fig. 2B). In most other cases, the gene was partially recoverable, but fragmentation of the assembly precluded assessment of functionality (Fig. 2B). Again, the lowest quality genome assemblies, *Lepraria sp. 1, S. cf. fronduliferum*, and *S. gregarium* had the most fragmented genes, 14, 12, and 10, respectively. Across the remaining genomes, eight other genes were incomplete. In two additional individual cases, *pro44* in *L. neozelandica* and *cas3* in the *S. virgatum* reference, we identified putative internal stop codons. However, these genes appeared functional in the remaining genome assemblies for both *Lepraria* and *Stereocaulon*. As with the meiosis genes above, RELAX analyses did not, with the exception of the COP9 signalosome genes, show an increased incidence of relaxation of selection within any functional category in the *Lepraria* lineage (Table 3).

**Table 3:** Selection on ascomata development genes in asexual *Lepraria* compared to sexual *Stereocaulon*.

| Gene | Group | n | K | p-value | Selection |
| --- | --- | --- | --- | --- | --- |
| *atg12* | Autophagy genes | 23 | 2.4 | 0.0299 | intensification |
| *atg4* | Autophagy genes | 26 | 0.43 | 0.1597 | none |
| *atg7* | Autophagy genes | 26 | ** | ** | ** |
| *atg8* | Autophagy genes | 26 | 0.58 | 0.1999 | none |
| *nbr1* | Autophagy genes | 24 | 0 | 0 | relaxation |
| *asf1* | Chromatin modifiers | 26 | 8.15 | 0.0031 | intensification |
| *cac2* | Chromatin modifiers | 26 | 0.05 | 0.0009 | relaxation |
| *crc1* | Chromatin modifiers | 26 | 4.1 | 0 | intensification |
| *rtt106* | Chromatin modifiers | 25 | 0.69 | 0.0069 | relaxation |
| *spt3* | Chromatin modifiers | 25 | 0.28 | 0.0241 | relaxation |
| *csnA* | COP9 signalosome | 25 | 0.58 | 0.0012 | relaxation |
| *csnB* | COP9 signalosome | 26 | 0.56 | 0 | relaxation |
| *csnC* | COP9 signalosome | 24 | 0.31 | 0.0097 | relaxation |
| *csnD* | COP9 signalosome | 25 | ** | ** | ** |
| *csnE* | COP9 signalosome | 25 | 0.53 | 0.0587 | none |
| *csnF* | COP9 signalosome | 25 | 0.45 | 0.0003 | relaxation |
| *csnG* | COP9 signalosome | 22 | 0.53 | 0.0282 | relaxation |
| *mak1* | MAPK signaling | 26 | 0.63 | 0.0006 | relaxation |
| *mek1* | MAPK signaling | 26 | 3.59 | 0.0001 | intensification |
| *mik1* | MAPK signaling | 25 | 11.64 | 0 | intensification |
| *pro40* | MAPK signaling | 24 | 1.8 | 0.0019 | intensification |
| *nor1* | NOX complex | 23 | 1.41 | 0.2202 | none |
| *nox1* | NOX complex | 24 | 6.2 | 0 | intensification |
| *pro41* | NOX complex | 26 | 1.68 | 0.0891 | none |
| *acl1* | Primary metabolism | 24 | 0 | 0.0002 | relaxation |
| *cas1-2* | Primary metabolism | 22 | 1.59 | 0.0389 | intensification |
| *cas3* | Primary metabolism | 24 | 0 | 0.0852 | none |
| *leu1* | Primary metabolism | 23 | 1.46 | 0.0857 | none |
| *sec22* | Primary metabolism | 26 | 1.35 | 0.0391 | intensification |
| *smac 06770* | Primary metabolism | 24 | 1.46 | 0.194 | none |
| *gsa1* | Signal transduction | 25 | 9.19 | 0 | intensification |
| *gsa2* | Signal transduction | 26 | 3.57 | 0.0271 | intensification |
| *gsa3* | Signal transduction | 26 | ** | ** | ** |
| *sac1* | Signal transduction | 27 | 0.43 | 0 | relaxation |
| *kin24* | Subunits of the STRIPAK complex | 25 | 6.47 | 0 | intensification |
| *kin3* | Subunits of the STRIPAK complex | 26 | 4.38 | 0 | intensification |
| *mob3* | Subunits of the STRIPAK complex | 25 | 1.08 | 0.7335 | none |
| *pp2ac1* | Subunits of the STRIPAK complex | 26 | 13.67 | 0.0022 | intensification |
| *pro11* | Subunits of the STRIPAK complex | 26 | 0.28 | 0 | relaxation |
| *pro22* | Subunits of the STRIPAK complex | 24 | 9.19 | 0 | intensification |
| *pro45* | Subunits of the STRIPAK complex | 26 | 8.3 | 0 | intensification |
| *mcm1* | Transcription factors | 26 | 10.18 | 0.0504 | none |
| *pro1* | Transcription factors | 26 | 0 | 0 | relaxation |
| *pro44* | Transcription factors | 25 | 19.97 | 0 | intensification |
| *ste12* | Transcription factors | 25 | 7.96 | 0 | intensification |

## Discussion

Our comparative genomic analyses demonstrate that, despite its apparent asexuality, *Lepraria* species likely maintain the capacity to reproduce sexually. The mating type locus appears complete and functional in all *Lepraria* species sampled. Additionally, genes in the meiosis toolkit and involved in the development of reproductive structures are present, likely functional, and not, in general, under relaxed selection in *Lepraria*, compared to the sister sexual clade *Stereocaulon*. Future work will probe why these genes remain functional in the absence of canonical sexual reproduction in these lichenized fungi.

### Lepraria is heterothallic and harbors both MAT idiomorphs

We identify intact mating loci and putatively functional *MAT* genes in all *Lepraria* and *Stereocaulon* genomes. In each individual, the *MAT* locus appears to be single-copy (i.e., only one *MAT* locus was recovered from each individual), suggesting that all sampled individuals are heterothallic. This is in line with expectations based on recent work, which did not include *Lepraria* or *Stereocaulon*, but found that the overwhelming majority of Lecanoromycetes genomes are heterothallic [22, 59, 69]; but, see White et al. [23] demonstrating homothallism in a few Teloschistaceae. Our results also confirm the findings of Pfeffer et al. [70] that the *L. neglecta* genome included a complete *MAT* locus with functional *MAT1-2* genes and a putative *MAT1-1* pseudogene, while extending the results on multiple fronts. First, we provide evidence of functional *MAT* loci in a further 11 *Lepraria* species (12 genomes; Fig. 1). Second, we demonstrate that the *Lepraria* genus is heterothallic and harbors functional *MAT1-1* and *MAT1-2*. If the entire *Lepraria* lineage had been asexual since early in its evolutionary history, one may expect a single *MAT* idiomorph to have fixed somewhere along the lineage, but we see both idiomorphs throughout the tree (Fig. 1). In fact, the two sampled *L. finkii* individuals are of opposite idiomorphs, suggesting the potential for ongoing sexual reproduction in this species, as well as the genus more broadly. Lastly, we add known functional *MAT* sequences from the obligately sexual sister genus *Stereocaulon* and show that the genes and structure of the *MAT1-1* and *MAT1-2* idiomorphs are largely conserved between the two genera (Fig. S3).

Our *MAT* loci are largely consistent with other fungi of the phylum Pezizomycotina (filamentous ascomycetes), where the *MAT* transcription factors are typically located between *apn2* and *sla2* [71, 72]. Within the locus, sequence synteny is consistent with closely related genera, including *Letharia* [59] and *Cladonia* [69]. Moreover, all identified accessory genes were located in identical positions as in other Lecanoromycetes, specifically between *MAT1-1-1* and *sla2,* and between *MAT1-2-1* and *apn2* [22]. The putative pseudogenized *MAT1-1-1* exon found downstream of our *Lepraria MAT1-2-x* accessory gene is consistent with many Lecanoromycetes, including Parmeliaceae (*Letharia*), Cladoniaceae (*Cladonia*), and Umbilicariaceae [59]. We show here that the architecture of both idiomorphs extends to Stereocaulaceae as well. However, this architecture differs in Collemataceae and Icmadophilaceae, so it is not conserved across all Lecanoromycetes [59].

While the presence and putative functionality of *MAT* loci suggest the possibility of sexual reproduction in *Lepraria*, it does not preclude alternative roles for this locus. For example, the basidiomycete fungus *Malassezia* has lost nearly all meiosis-specific genes (see “*Lepraria maintains genes required for meiosis*” below), but maintains an apparently functional *MAT* system [13, 73]. Similarly, a mating type locus has been identified in the genome of the multinucleate arbuscular endomycorrhizal fungus (AMF) *Rhizophagus irregularis*, a so-called “ancient asexual” [74, 75]. Studies even suggest that the coexistence of different mating types in a cell leads to the expression of genes involved in mating and meiosis [76, 77]. However, the dearth of evidence for sexual structures in *Lepraria* suggests a parasexual process may be more likely than sexual reproduction.

### Lepraria maintains genes required for meiosis

We were able to identify complete annotations for all nine core meiosis genes and 20 associated mitosis genes in our *L. finkii* reference genome. In large part, these genes had conserved start codons and introns, and relatively consistent stop codons among 13 *Lepraria* and 11 *Stereocaulon*, as well as the two *Cladonia* outgroup species. Exceptions included *rad50*, for which we could not identify a conserved start codon; however, the start of *rad50* is also not conserved in other systems. Additionally, it is counterintuitive that *hop2* appears functional in asexual *Lepraria*, but seems to be undergoing pseudogenization in sexual *Stereocaulon*. However, Sordariomycete fungi, including the sexual *Neurospora crassa,* complete meiosis without orthologs of *mnd1*, *hop1*, or *hop2* [78].

If *Lepraria* has indeed lost the functions of meiosis and sexual reproduction, we would expect to observe a general relaxation of selection across the genes involved [79], which we did not. Many of the genes we examined have secondary functions, which explain the maintenance of selection [61–63]; however, this is less likely for the nine meiosis-specific genes (*dmc1*, *hop1*, *hop2*, *mer3*, *mnd1*, *msh4*, *msh5*, *rec8*, and *spo11*). We do see a slight relaxation of selection in *mer3* and *spo11*, and significant relaxation in *mnd1*, although Sordariomycetes complete their sexual cycles without this gene [78]. For the remaining six genes, it is plausible that an insufficient amount of time has passed since *Lepraria* became asexual to accumulate mutations that would reveal a pattern of relaxed selection. However, the estimated divergence time between *Lepraria* and *Stereocaulon* is at least 30M years [80], which should be sufficient to observe relaxed selection, pseudogenization, or even gene loss [79]. Thus, the lack of relaxation across meiosis-specific genes suggests continued use. Meiotic proteins could have evolved new functions in the asexual *Lepraria* lineages; however, two primary lines of evidence argue against this. First, it is highly unlikely that neofunctionalization would have occurred in all six meiosis-specific genes that do not show evidence of relaxed selection (*dmc1*, *hop1*, *hop2*, *msh4*, *msh5*, and *rec8*). Second, with neofunctionalization, we would expect *Lepraria* and *Stereocaulon* genes to have diverged substantially, but they remain highly conserved (Fig. S4).

In general, bioinformatic identification of meiosis genes in putatively asexual lineages has revealed that most of these organisms have the tools necessary to complete meiosis [13]. Exceptions include loss of the complete complement in nucleomorphs and the majority of meiosis-specific genes in *Malassezia*, while bdelloid rotifers have lost most genes involved in recombination and crossover resolution [13]. Much more common are cases such as the basidiomycete fungus *Pseudozyma* [16] and AMF *Glomus* [11], where genomes are found to contain complete and seemingly functional complements of core meiotic genes. Presence does not necessarily prove use in meiosis, but evidence for recombination has been found in AMF genera *Glomus* and *Rhizophagus* [81, 82].

### Genes for fruiting body development remain in Lepraria

Despite the complete absence of ascomata development across the *Lepraria* lineage, most genes identified as crucial to the development of fungal sexual reproductive structures [62, 63] appear to be present and functional. We interpret these findings with caution for several reasons. First, although the model fungi in which these genes were examined are also Ascomycota, they are highly diverged from lichenized fungi. Thus, we cannot be sure that the functions of these genes are conserved. Additionally, the pleiotropic effects of many of these genes make it difficult to assume that presence and potential functionality indicate the ability to form ascomata; many likely have other essential functions and remain in use and under selection.

The COP9 signalosome is the only instance where we found that the majority of genes show evidence of mildly relaxed selection in the *Lepraria* lineage. Although this complex is important in post-translational processes across Eukaryotes, it does not appear to be essential for fungal viability [83]. Work on the filamentous fungus *Aspergillus nidulans* has demonstrated a critical role for most components of the COP9 signalosome involved in timing the sexual cycle in response to light [62, 83]. Knocking out *csnA*, *csnB*, *csnD*, or *csnE* prevents the formation of proto-ascomata structures [84, 85]. Perhaps the COP9 signalosome lacks another function in *Lepraria*, and thus its components are no longer under selection.

### Potential for parasexual reproduction in Lepraria

Despite the maintenance of the machinery for mating, meiosis, and the formation of sexual reproductive structures, the fact remains that across 200 years of careful observation, canonical sexual reproduction (i.e., the formation of ascomata and meiotic generation of ascospores) has never been observed in *Lepraria* [25, 86, 87]. In many cases of non-lichenized “*fungi imperfecti*,” sexual reproduction was eventually observed on rare occasions or under specialized conditions (reviewed in [88]), but the consensus among lichenologists is that this is extremely unlikely to be the case for *Lepraria* since it forms long-lived vegetative structures and the ascomata in the sister-group *Stereocaulon* are visible for several years [30]. The possibility remains that *Lepraria* engages in a form of parasexual reproduction that utilizes mating type and core meiotic genes. For example, the asexual yeast *Candida albicans* completes a parasexual cycle that involves both. Conjugation occurs between a and ɑ mating strains [89] and the meiosis-specific *spo11* has been co-opted for mitotic recombination between homologous chromosomes [90]. Tripp and Lendemer [18] provided evidence from *L. pacifica* and *L. squamatica* to suggest that fusion can occur between hyphae of two individuals, two soredia, or soredia and hyphae, but did not resolve whether or not the individuals involved may be genetically distinct. They also suggest that fusion could lead to diploidy, with subsequent parasexual mitotic recombination and random loss of chromosomes to regain a haploid state. If this is the case, either fusion or initiation of recombination could require opposing *MAT* idiomorphs and mitotic recombination may very well mirror meiosis and involve putatively meiosis-specific genes. This could explain “neofunctionalization” across meiosis genes in *Lepraria* without substantial divergence from *Stereocaulon*. Future population-level genomic data will be needed to document the frequencies of recombination and *MAT* idiomorphs across *Lepraria* species.

## Conclusions

Recently Hofstatter and Lahr [13] proposed a paradigm shift: “All eukaryotes are sexual, unless proven otherwise”. Bioinformatically, we show that *Lepraria* has the capacity for sexual reproduction, but should we assume it is sexual? Asexual reproduction has many benefits, which are all the more heightened by the symbiotic relationship central to lichenized fungi. In lichens, clonal reproduction via soredia (isidia, etc.) is unique and has the outsized advantage of maintaining an unbroken connection between the fungus and its algal symbiont. Thus, a sort of obligate parasexuality may be particularly advantageous in lichens to maintain the benefits of asexuality (i.e., retaining its symbiont) without giving up the short and long-term benefits of recombination and genome maintenance, respectively. In other fungi, mitotic recombination has already been shown to accelerate adaptation [91], which could help explain why speciation seems as common in *Lepraria* as in sexual lichens. *Lepraria* species are diversified chemically and morphologically, and they maintain distinct species across broad distributions, something usually thought to involve sexuality [25, 28]. Furthermore, an obligate parasexuality of *Lepraria* explains how it has survived without sex for over 30M years [80]. Perhaps the *Lepraria* lineage adopted an alternative life history strategy as it diverged from *Stereocaulon*. Where *Stereocaulon* maintained obligate sexual development, *Lepraria* instead goes “all in” on the production of soredia and uses meiosis machinery for mitotic recombination instead, reaping the benefits of asexuality without paying the costs.

## Declarations

### Competing interests

The authors declare that they have no competing interests.

### Funding

This research was funded by the Grainger Foundation and the Negaunee Foundation. Felix Grewe’s fieldwork on Livingston Island was supported by CRYPTOCOVER (Spanish Ministry of Science CTM2015-64728-C2-1-R), headed by Leopold Garcia Sancho.

### Authors’ contributions

FG and HTL designed the project. MMD, YS, and ABP collected and analyzed the data. MMD drafted the manuscript. All authors provided edits and approved the final manuscript.

## Acknowledgments

The authors thank the University of Wisconsin Biotechnology Center DNA Sequencing Facility and the Rush Genomics and Microbiome Core Facility for providing next-generation sequencing consultation and services. From The Field Museum of Natural History, we thank the Pritzker Laboratory for Molecular Systematics and Evolution for DNA extraction services and the Grainger Bioinformatics Center for computational resources. We extend our gratitude to the following individuals for their invaluable assistance with collections: Gintaras Kantvilas in Australia; Juan Larrain and Matt von Konrat in Chile; Peter De Lange, Jeremy Rolf, Dan Blanchon, Allison Knight, and John Knight in New Zealand; and Ulrike Ruprecht, Christian Printzen, Leopold Garcia Sancho, and Ulrik Søchting in Antarctica. We also thank the staff of the Spanish Antarctic Station Juan Carlos I. Additionally, we want to thank Todd Widhelm for his collection support and help with the Field Museum herbarium. We also appreciate the technical support provided by Kevin Feldheim at the Pritzker Laboratory at the Field Museum, and contributions from Isabel Distefano. Our thanks extend to Allison Knight, Ulrik Søchting, and Joel Mercado for providing samples, and the Chicago Botanic Garden for allowing us to collect *Lepraria* in their garden.

## Notes

### Competing Interest Statement

The authors have declared no competing interest.

