## Supplemental Figures for "Rethinking asexuality: the enigmatic case of functional sexual genes in *Lepraria* (Stereocaulaceae)"

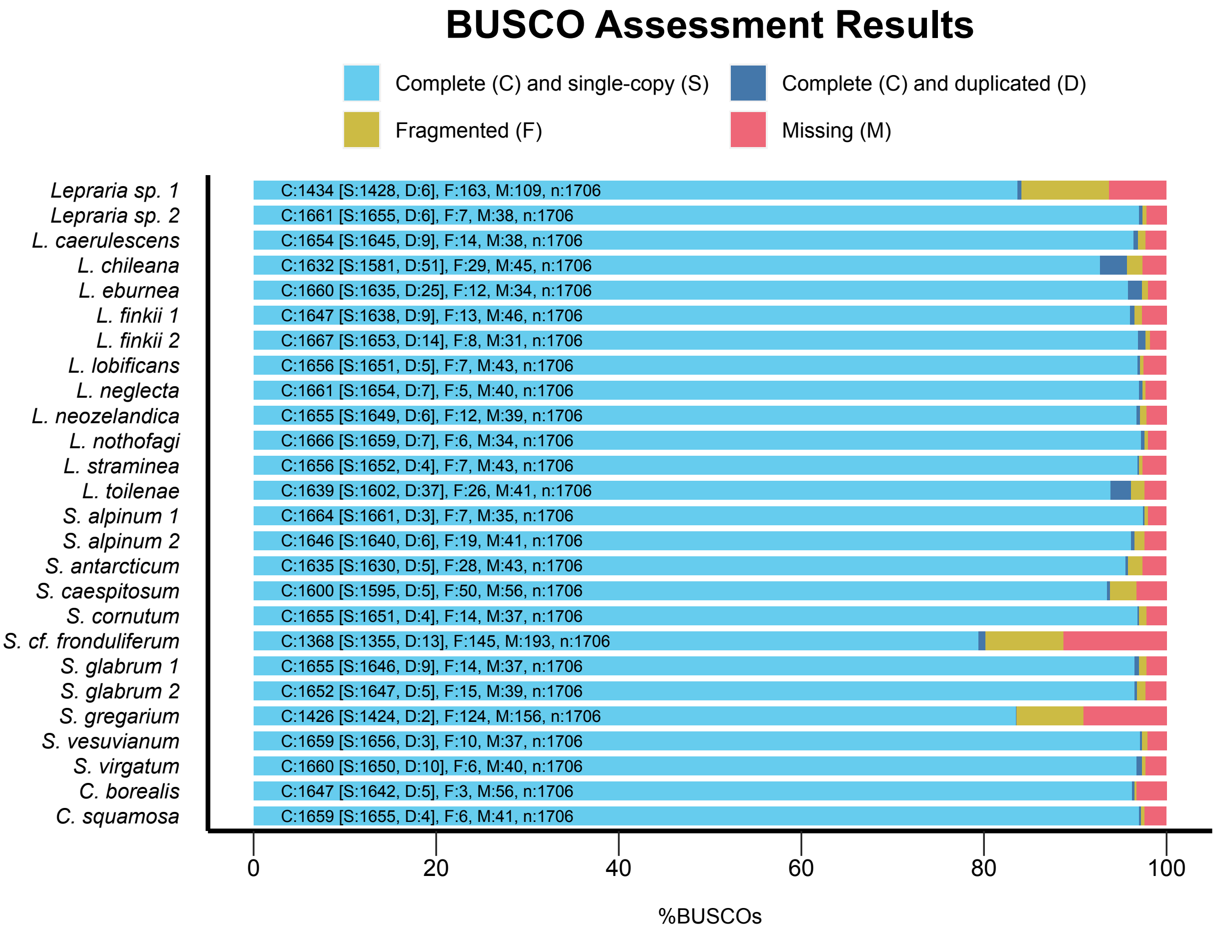


**Figure S1:** BUSCO assessment results for the 26 genomes included in our study, showing the percentage and category of single-copy orthologs from the Ascomycota data set (total n=1706) in each genome assembly.

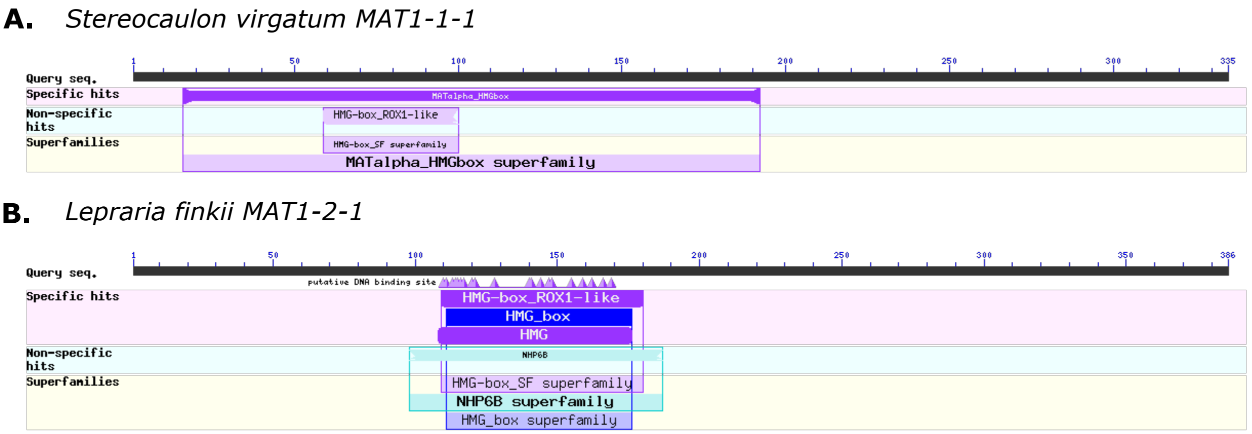


**Figure S2:** Results of *blastp* searches for conserved domains in *Stereocaulon* and *Lepraria MAT* genes. A) The *S. virgatum MAT1-1-1* gene contained the expected alpha domain. B) A high mobility group (HMG) domain was identified in the *L. finkii MAT1-2-1* gene.

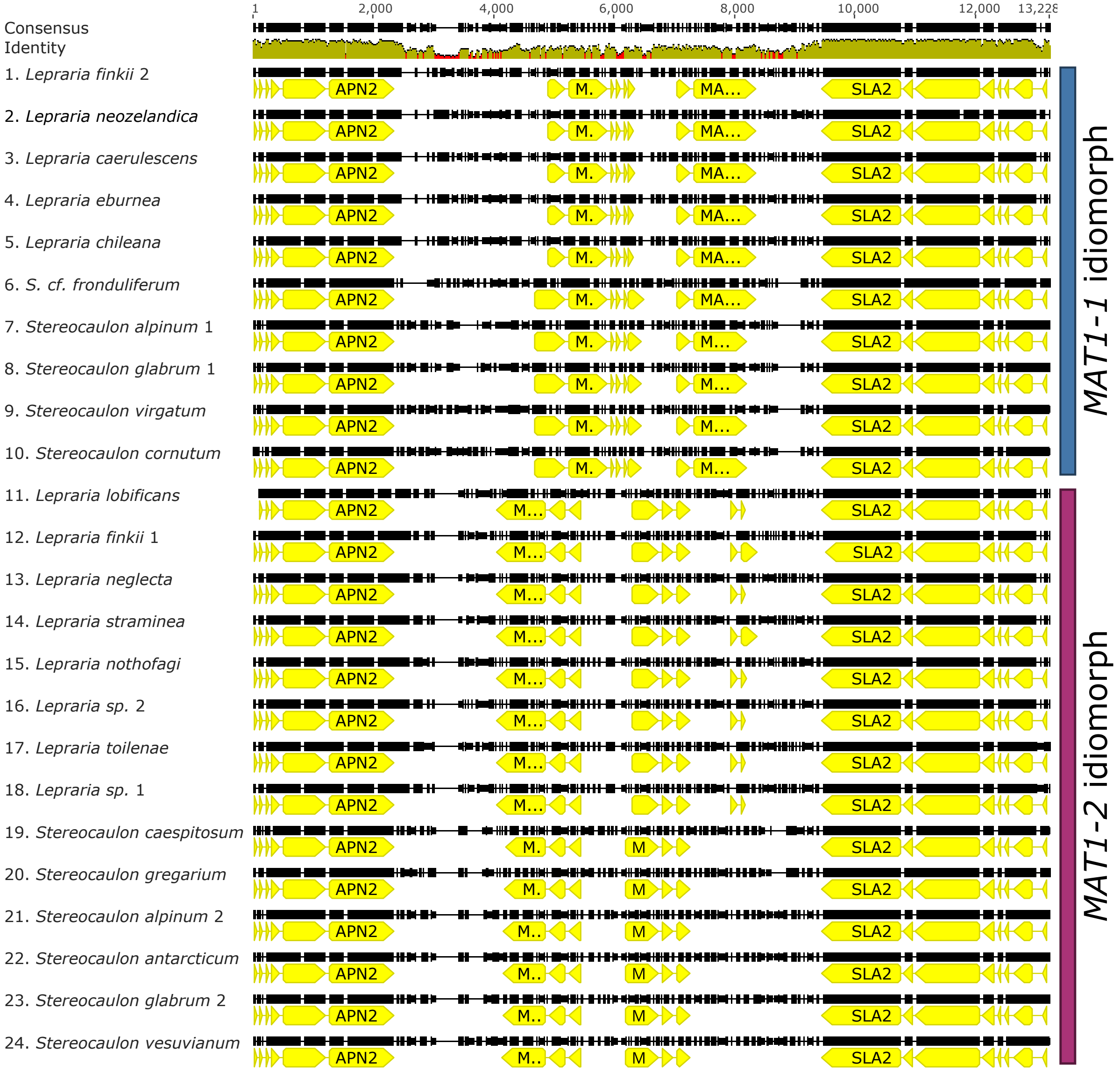


**Figure S3:** A Clustal alignment of all 24 annotated *Stereocaulon* and *Lepraria MAT* loci. See Fig. 1A for names of genes contained within the *MAT1-1* and *MAT1-2* idiomorphs. The figure was generated with Geneious version 2023.1 created by Biomatters (available from https://www.geneious.com).

**A.** *Stereocaulon virgatum dmc1*

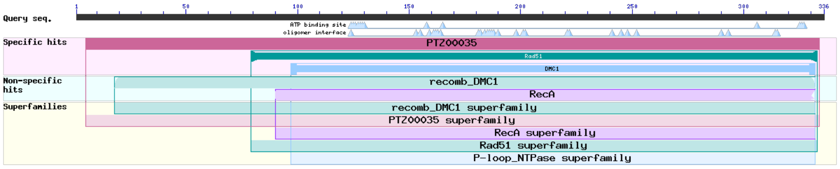


*Lepraria finkii dmc1*

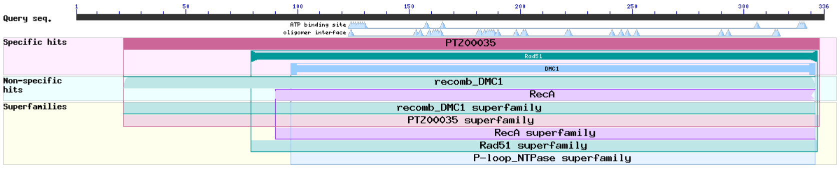


**B.** *Stereocaulon virgatum hop1*

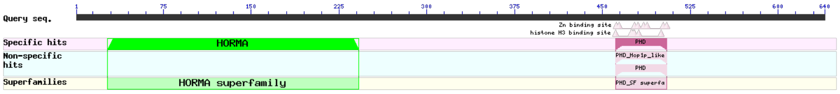


*Lepraria finkii hop1*

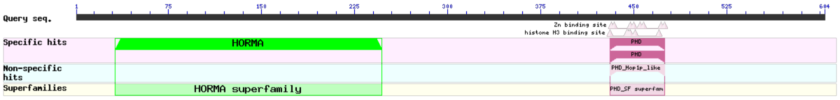


**C.** *Stereocaulon virgatum hop2*

No conserved domains found

*Lepraria finkii hop2*

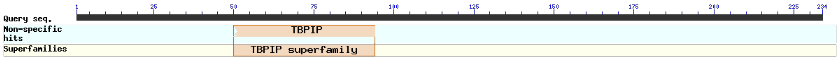


**D.** *Stereocaulon virgatum mnd1*

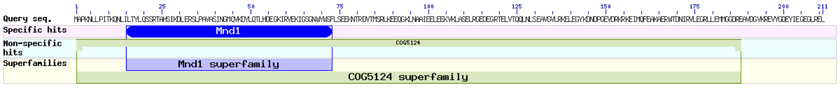


*Lepraria finkii mnd1*

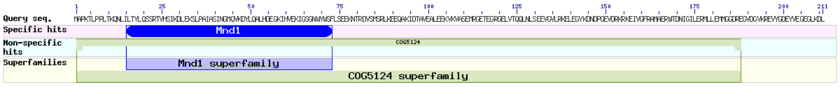


**E.** *Stereocaulon virgatum spo11*

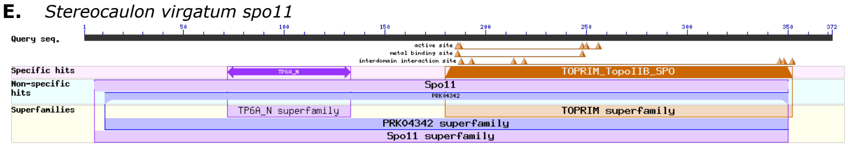


*Lepraria finkii spo11*

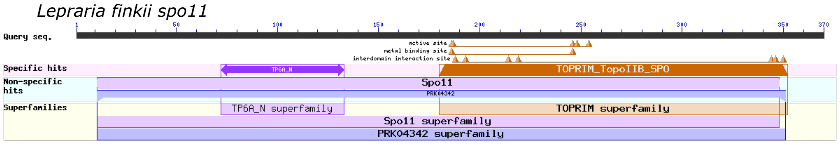


**F.** *Stereocaulon virgatum mer3*

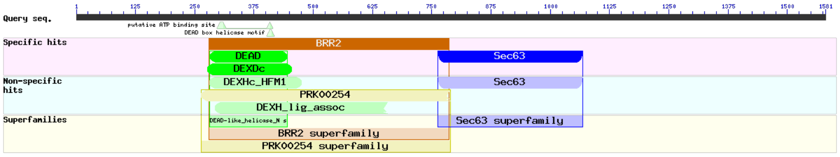


*Lepraria finkii mer3*

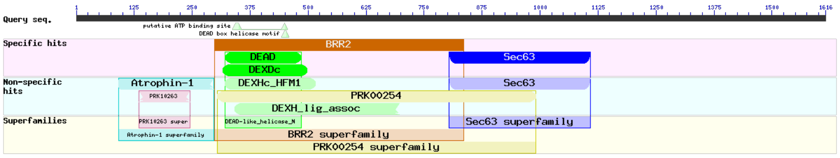

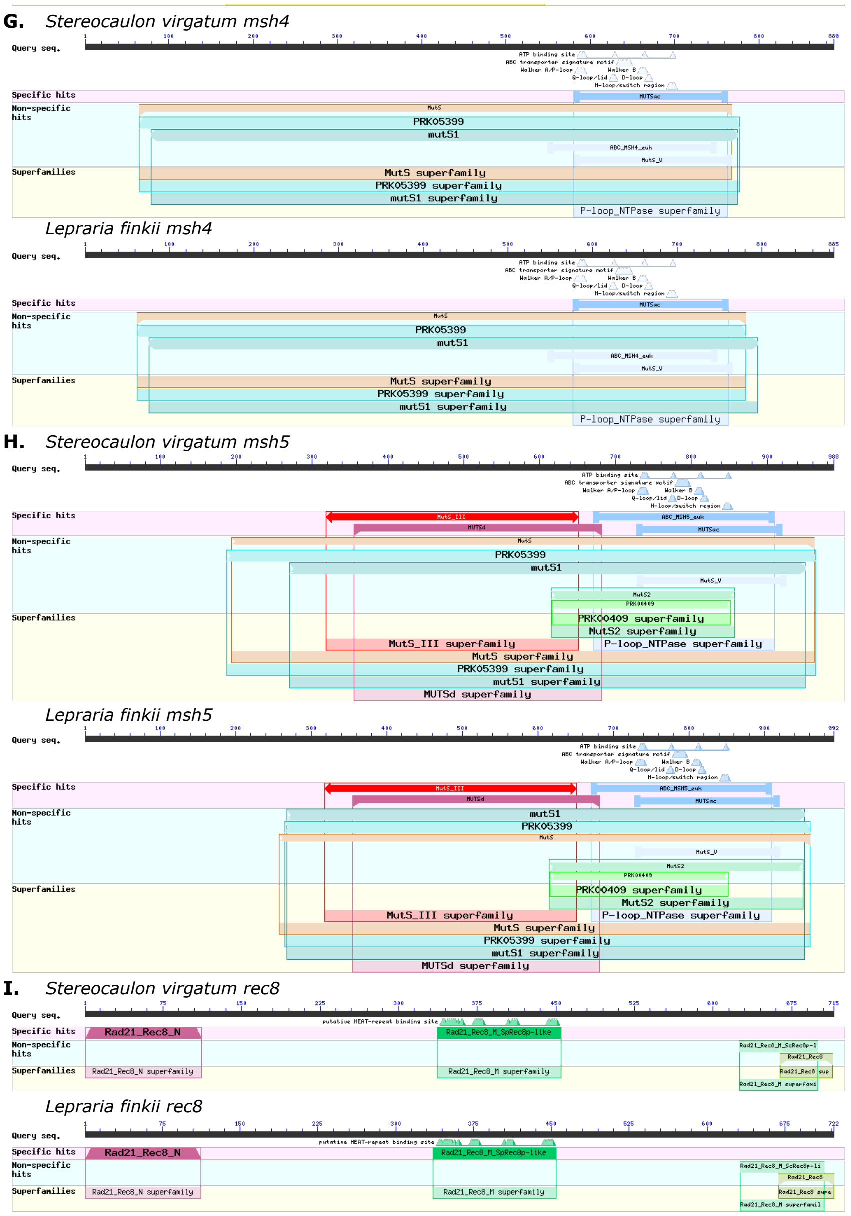


**Figure S4:** A compilation of *blastp* searches for conserved domains in the nine meiosis-specific genes for *Stereocaulon* and *Lepraria*, respectively.
